# Co_3_O_4_/PAn Magnetic Nanoparticle-Modified Electrochemical Immunosensor for Azocyclotin

**DOI:** 10.1101/2021.03.22.436409

**Authors:** Weihua Wang, Zhanjiang Han, Qian Xi, Dongla Gao, Ruicheng Guo, Pengcheng Kuang, Dongliang Li

## Abstract

In the current study, for rapid detection of azocylotin residues in agricultural products, the Co_3_O_4_/PAn nanoparticles modified electrochemical immunosensor was generated successfully. Azocylotin-BSA artificial antigen coupling to the surface of the working electrode coated with the Co_3_O_4_/PAn nanoparticles thin-layer, the competitive inhibition reaction is launched between the azocylotin in the samples and the azocylotin coupled on the electrode with the azocylotin monoclonal antibodies in the test system. The antigen-antibody reaction signal conductive amplified by the coupled silver nanoparticles, and then the electrolytic current in the reaction system was detected. After establishing basic detection system, a series of optimization including the concentration of immobilized membrane optimization, electrode surface coated material composition optimization, selection of buffer, coupling antigen concentration optimization and anti-antibody label optimization will be done. Subsequently, the concentrations of azocylotin in samples of standards, orange and apple were tested, and the results indicated that this immune sensor has good sensitivity and high accuracy. The research could provide the reference for the research, development and application of sensors to realize rapid detection of pesticide residues in agricultural products.

## Introduction

The tri(cyclohexyl)-1H-1,2,4-triazole-1-yltin (azocylotin), one kind of organometallic contact acaricide of triazole with moderate toxicity, was widely used in killing mites of crops, for the good efficacy in controlling nymph, adult mite and summer egg [1-3].

Currently, in order to detect the content of the residual azocylotin in fruit, vegetable and soil, both gas chromatography (GC) and liquid chromatography (LC) are mainly used. LC with UV detection allows to directly detecting azocylotin and cyhexatin of food but has a lower sensitivity. Moreover, these two agents are easily infected by impurities at the wavelength of 220 nm and tend to be decomposed in case of exposure to UV-light [4,5]. GC can only detect the azocylotin of the samples which are processed by a derivatization method, so this method brings a relatively complex procedure of detection [6,7].In recent years, some immunoassay methods based on antigen-antibody reaction are being used gradually to detect the pesticide residue in food and environment, and they are replacing gradually mass spectrometry and chromatography in the leading position to detect pesticide residues [8,9].Due to high sensitivity, strong specificity, economic applicability and adaptability in developing the rapid on-site detection product.

Food safety not only matter the human health, but also become a major factor affecting the market competitiveness of agricultural products in international trade, with the progression of globalization and agricultural industrialization development. These have become the most effective technological means for protection of agricultural products and food trade in many countries. For example, Japan executed Positive List System in May 29, 2006, which proposed a limit standard for the majority of pesticides, veterinary drugs and agricultural chemicals (e.g., as per the regulation, azocyclotin and cyhexatin must not be detected in food, and the limit of detection should be 0.02mg/kg [7]. In order to adapt to the international development trend, enhancing the export competitiveness of agricultural products in our country and guarantee the food safety of the domestic consumers and establishing a rapid and effective detection method of the pesticide residues in agricultural products has becoming extremely urgent.

Generally, Nano material which is considered as the most promising material in 21 century, for the granule and surface effect, shows many distinct physical and chemical characteristics and has a bright application prospect in chemistry, biology, medicine, aerospace and environment monitoring. Especially, the nano technology has played a great role in designing or producing antibacterial materials, drug carriers, biosensors and environment monitoring [11]. The concentration of the sample to be tested could be determined by measuring the changes of the current and conductivity between reference electrode and working electrode based on the electrochemical reaction between electrode and the sample to be tested, using the electrochemical sensor, which contained the electrode modified and prepared by metal/metallic oxide-derived nano materials.

As an important metal oxide ceramic material, Co_3_O_4_ is widely used in electrochemistry, magnetics and catalysis field, and the application of Co_3_O_4_ in the super capacitor has become a hot spot [12,13]. In the practical application, the micro structures of Co_3_O_4_ such as grain size and crystal morphology, are the key parameters to determine its performance [14]. In the current study, based on the strong and selective adsorption of Co_3_O_4_ nanocomposites to phosphate group, a self-made working electrode was adopted, which was assembled with Co_3_O_4_/PAn magnetic nanoparticles, and the voltammetry was applied to measure azocylotin concentration of samples. Hence, building a sensitive, rapid, cheap and simple detection method for triazole pesticide residue was preliminarily created. Here we report the preparation process of the electrodes and the detection method of azocylotin.

## Materials and Methods

### Generation of Co_3_O_4_ nanoparticles

A chemical coprecipitation method was used to generate Co_3_O_4_ nanoparticles. The detailed process was as follows: firstly, 0.2 g Co(NO_3_)_2_ was dissolved successfully in 5 ml alcohol and 10 ml distilled water by magnetic stirring; secondly, 2.5 ml o ammonia water was added dropwise into the mixture above and adjusted to pH=8-9 by Sodium hydroxide The chemical reaction (Fig. 1 A) happened under magnetic stirring, and the purplish red mixer became black. The black solution was filtered with an air pump, and then washed repeatedly. The product of filtration was dried in an oven at 60°C, and Co_3_O_4_ nanoparticles were obtained finally.

**Fig 1.**
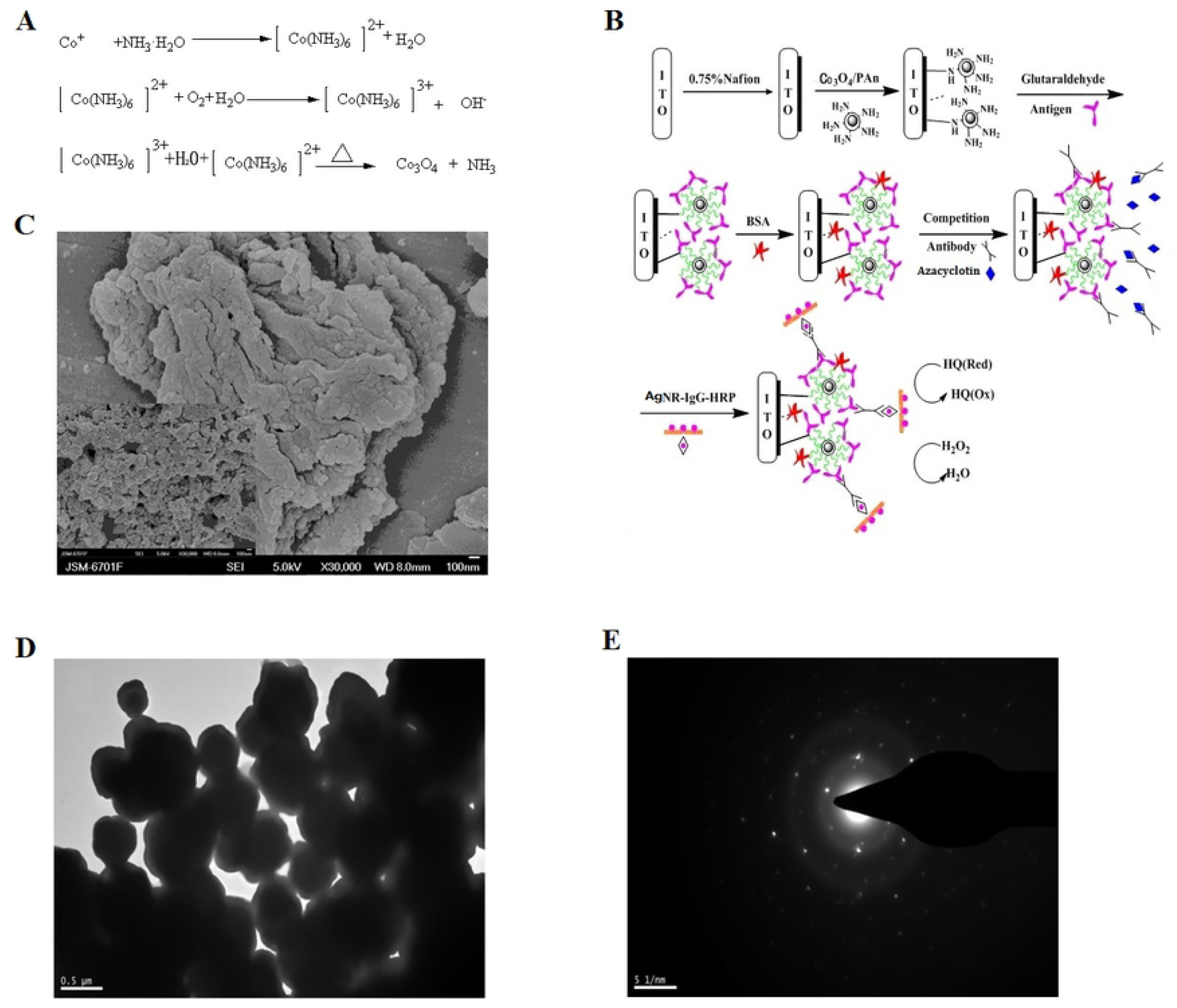
The generation of azocyclotin immunosensor. (A) Synthetic route of Co_3_O_4_ nanoparticle; (B) Determination principle of azocyclotin immunosensor; (C) Scanning Electron Microscope Image of Co_3_O_4_/PAn nanocomposites; (D) Determination of silver nanorods morphology and size, TEM image of nanometric copper rod; (E) Electronogram of nanometric copper rod.

### Generation of core-shell magnetic nanoparticles (Co_3_O_4_/PAn)

0.36 g CTAB was dissolved in 50 ml distilled water, and a certain amount of Co_3_O_4_ nanoparticles were then added. After stirring the mixture for 5 h, 180 μl aniline was added, and the chemical reaction lasted for 1 h. Subsequently, 0.6g citric acid was added into the mixture above, and the chemical reaction lasted for 10 h with stiring. At last, post addition of 0.7g ammonium persulfate, the chemical reaction lasted for 12 h with stiring, and then was terminated. The resulted solution was filtered with a vacuum pump, and the filtrate was retained. The filtrate was washed repeatedly with deionized water and ethyl alcohol until the filtrate was colorless, and the Co_3_O_4_/PAn was contained in the final filtrate. The solid samples could be obtained by vacuum drying at 80 °C.

### Generation of silver nanoparticles

0.1 ml mM AgNO_3_ was diluted to 1 ml, and then was mixed with 5 ml 0.2 M CTAB, Subsequently, 0.1g NaBH_4_ was added into the mixer above, and magnetic stirring lasted for 2 min at 30°C. At the last, the mixer was kept ageing for 5 h in a water bath at 28 °C, and the seed solution was generated successfully.

Afterwards, 5 ml 0.2 M CTAB was put into a big beaker, 15 ml mM AgNO_3_ was put into three small breaker averagely with shaking for one min, and 100 μl 0.1M ascorbic acid was added into each breaker. The chemical reaction lasted for two hours with magnetic stirring, and then 60 μl 0.01 M AgNO_3_was added into each baker. These breakers were kept in a thermostatic water bath at 28 °C after shaking, and the growth solution was obtained successfully.

The growth solution was kept in a thermostatic water bath for 10 min, and 5 ml seed solution was added. The mixer above was shook for 5 min, and then kept in a water bath at 28 °C for 5 min. At the last, the silver nanoparticle solution was generated successfully.

### Enzyme-labeled second antibody labeled with silver nanoparticles

5 ml 0.05g/ml silver nanoparticle solution was adjusted to pH=8 with 0.1M NaOH solution. Subsequently, add 20 ml IgG-HRP and 20 ml 1 mg/ml HRP was added into the mixer above. After stirring for 30 min, the majority of IgG-HRP antibodies could be absorbed and presented on the surface of silver nanoparticles, and then centrifugation was implemented at 10,000 rpm for 20 min, so as to remove the uncombined enzyme-labeled second antibodies. The resultant should be washed repeatedly with PBS (0.01M, pH=7.4), and the obtained AgNRs-IgG-HRP was dispersed in 1 ml PBS buffer, which should be stored at 4 °C for later use.

### Generation of immunoelectrode

The ITO conductive glasses were cut into 2×4 cm slices. These slices were washed ultrasonically with acetone and ethyl alcohol and dried. Probes were sealed with the insulation paste with 1 cm^2^ left and then washed ultrasonicly using methylbenzene, acetone, ethyl alcohol and deionized water for several min, respectively. These slices should be dried with N_2_, then 100 μl Nafion solution (w/v) added on the working area. After the drying, 100 μl diluted Co_3_O_4_/Pan nanocomposites solution was dropped on the surface of ITO electrode, which was dried in an oven, and then 100 μl 0.25% glutaraldehyde solution added for activation at room temperature. After 2 h, the activated ITO electrode was washed with double-distilled water and dried in N_2_. Subsequently, 100 μl diluted azocylotin artificial antigen was added and incubated at 37 °C for 3 h. The electrode was washed with PBS repeatedly, and after drying, 100 μl 5 mg/ml BSA buffer for sealing in order to eliminate nonspecific adsorption. The electrode was kept at 37 °C for 5 h, and then washed repeatedly. Finally, the prepared sensor probe was stored at 4 °C for later use.

### Electrochemical detection

In order to detect azocylotin, an indirect competitive immunoassay was applied. To the close electrodes, 100 μl mixed solution, which contained azocylotin monoclonal antibodies and standard azocylotin, was dropped onto the electrode closed, and incubated at 37 °C for 1 h. Subsequently, the electrode was rinsed repeatedly with PBS buffer, and then 100 μl nanometric copper rod-labeled IgG-HRP antibodies dropped onto it with incubation at 37°C for 45 min. The rinsing procedure should be implemented for several times.

Finally, a three-electrode system, which is composed of the generated immunoelectrodes above, Ag/AgCl electrode and platinum electrode, was placed in a 10 ml beaker that contained 4 ml mM potassium ferricyanide mixture, a cyclic voltammetry scan was conducted at the scanning rate of 50 mV/s in a range of -0.6 to 0.8 V. The mixer should be stirred constantly with 10 μl 0.2M H_2_O_2_ being added slowly, and this procedure was repeated four times. With the increase of the concentration of H2O2, the cathode current declined continuously. Three parallel experiments were set in the same experiment, and the mean value of the results from the three experiments was taken. The determination principle was shown in Fig. 1B.

## Results

### FESEM characterization of Co_3_O_4_/PAn nanocomposites

The Co_3_O_4_/PAn nanocomposites were prepared, and their electron microscope images (FESEM) were scanned. It showed that Co_3_O_4_/PAn nanocomposites were mainly present in the form of nanoparticles, and some about 80 nm particles in size were connected and accumulated fluffily each other with some local agglomeration (Fig.1C). The generated Co_3_O_4_ was granular with diameter about 40 nm, loose accumulation and good dispersion. FESEM test showed that Co_3_O_4_ and polyaniline formed constitutionally stable composite, and this Co_3_O_4_/PAn nanocomposite might be good for improving the conduction ability of the material and the dynamic performance of the electrode process.

### Morphology and size of silver nanorods

The TEM images of silver nanoparticles were generated (Fig.1D & E) showing that the ascorbic acid-reduced silver nanoparticles were present in the form of nanoparticles, with diameter about 350 nm and good dispersion (Fig.1D). Moreover, it also indicated that these silver nanoparticles had the perfect crystallinity, were very beneficial for to connecting and labeling with the antibodies.

### Analysis of electrochemical behavior in the electrode assembly process

The Nafion solution was dropped onto the pre-treated ITO electrode, and formed a layer of immobilized membrane applied to adsorb Co_3_O_4_/PAn nanoparticles. The Co_3_O_4_/PAn nanocomposites were assembled onto the electrode, and then azocyclotin-BSA was covalently linked to the electrode by using glutaraldehyde as the cross-linking agent. Because the carboxyl on azocyclotin-BSA artificial antigen was able to have an amidation reaction with the amino group on the outer layer of activated Co_3_O_4_/PAn nanoparticles, therefore azocyclotin monoclonal antibodies could specifically bind to azocyclotin-BSA.

The electrochemical property of the electrode during self-assembly processes was investigated through cyclic voltammetry, and the behaviors of cyclic voltammetry of the immunosensor at different assembly phases and in 4 ml potassium ferricyanide solution were discovered (Fig.2). As shown in Fig.2, the curve a represents the cyclic voltammogram of redox current of the bare ITO electrode while the curve b indicated that when, the current of the cyclic voltammetry curve would decline if the electrode was modified with Nafion membrane, suggesting that Nafion membrane could hinder the electron transfer. The curve c revealed that the redox peak current of the modified electrode in the potassium ferricyanide solution could increase further after ion exchange occurred between Co_3_O_4_/PAn and Nafion membrane, indicating that Co_3_O_4_/PAn in Nafion membrane can effectively transfer the electrons, and the curve d revealed that the redox peak declined compared with curve c after the electrode was re-modified and coupled with azocylotin BSA, which impede electron transport. In particular, azocylotin artificial antigen specifically binds to azocylotin monoclonal antibody and forms the antigen-antibody complex, which can block more channels on the surface of modified electrode and increase the resistance across the membrane, causing further decrease of the response current (curve e).

**Fig 2.**
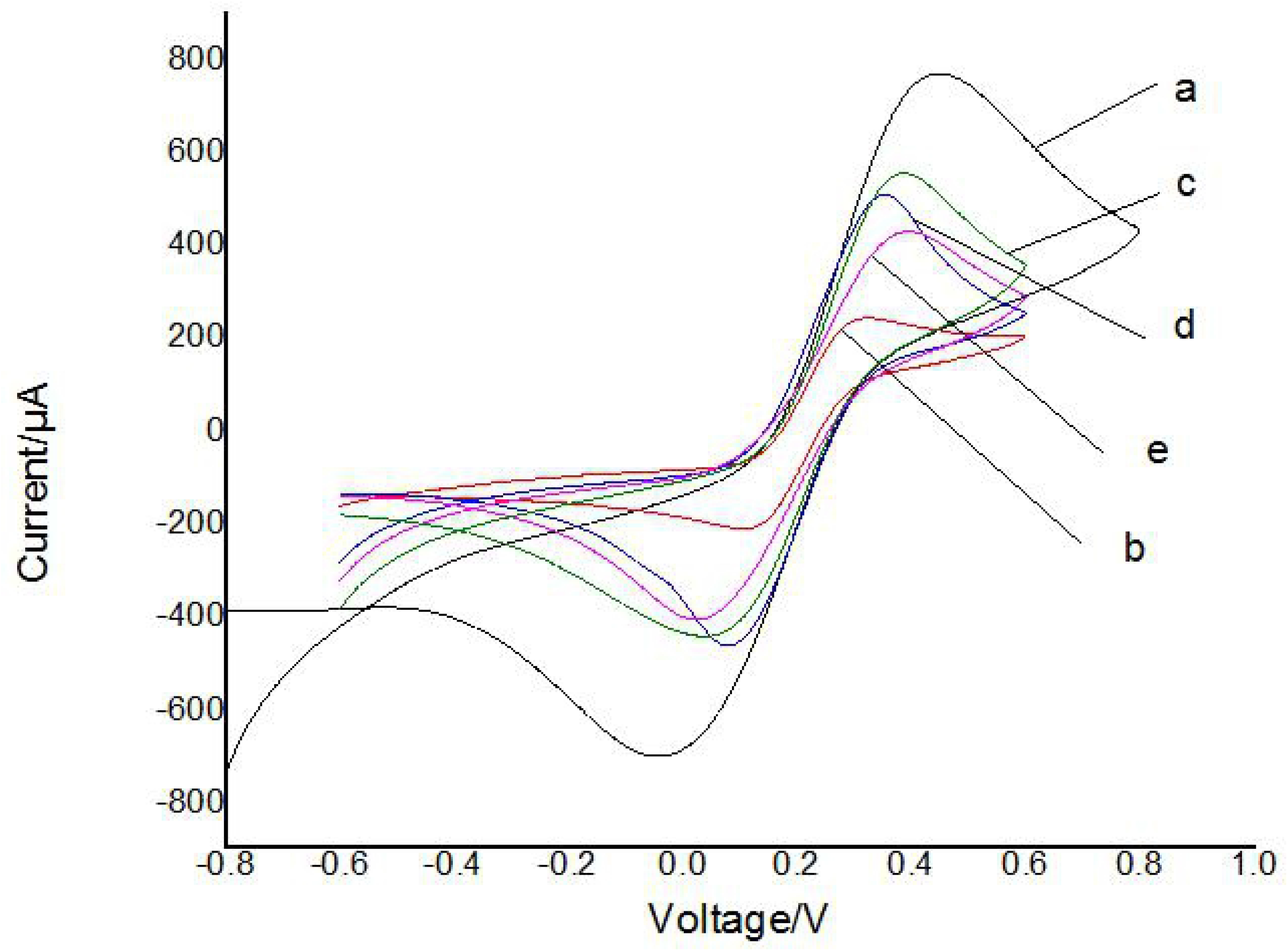
Analysis of electrochemical properties of the electrodes in the self-assembly processes. The curves (a, b, c, d and e) represent the bare, Nafion-modified, nano-Co_3_O_4_/PAn-modified azocyclotin artificial antigen-modified and azocyclotin monoclonal antibody-modified electrodes, respectively.

### Optimization of reaction conditions

It is very important to optimize conditions in the process of immunization for performance of the immunosensors. Generally, dilution ratio of the antigen buffer for immobilization, dilution ratio of the antibody buffer binding to antigen, the amount of antibody binding to hapten, the duration of immunoreaction and temperature have great impacts on the accuracy and sensitivity of the sensors. The experiment focused on optimizing concentration of Nafion solution concentration, dilution ratio of Co_3_O_4_/Pan solution, dilution ratio of antigen and antibody and pH value of diluted potassium ferricyanide buffer. In the optimization process, three parallel sub experiments were set in the same experiments, and the mean value of the results from the three experiments was taken.

### Optimization of immobilized membrane concentration

As shown in Fig. 3A, with the increase of Nafion concentration, the corresponding variation of electric current increased first and then decreases. It indicated that the increase of Nafion concentration could couple more Co_3_O_4_/Pan nanoparticles, but when the Nafion concentration exceeded 0.75%, the Nafion membrane on the ITO surface was too thick and not suitable for electronic transmission. Therefore, 0.75% was chosen as the optimum concentration for immunosensors in this experiment.

**Fig 3.**
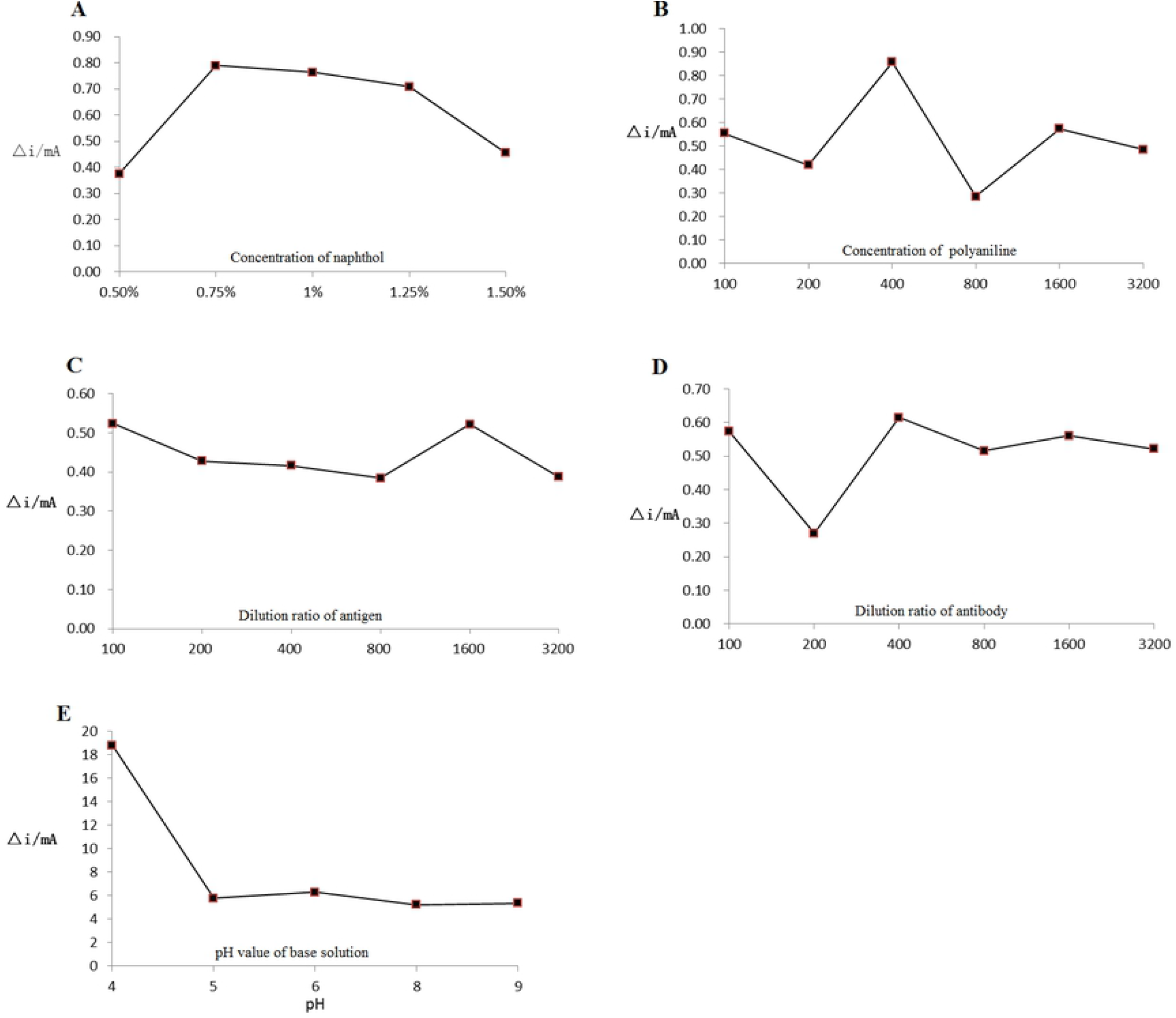
Optimization of conditions for performance of the immunosensors. (A) Optimization of naphthol concentration; (B) Optimization of polyaniline concentration; (C) Optimization of dilution ratio of antigen; (D) Optimization of dilution ratio of antibody; (E) Optimization of pH value of base solution.

### Selection of core-shell Co_3_O_4_/PAn nanocomposite concentration

The immobilization of antigen in the late experiment of the immunosensors was determined by the concentration of nanoparticle. Influences of different concentration ratios of core-shell nanoparticles on the immunosensors were investigated (Fig. 3B). The change of electric current had a certain fluctuation in the dilution ratio of Co_3_O_4_/PAn nanocomposites from 1:100 to 1: 400, and the maximum change appeared when the dilution ratio was up to 1:400. Moreover, there was little change if dilution ratio was over 1:400, indicating that with dilution ratio being 1:400, the binding force between Nifion and polyaniline nanoparticles was the strongest, showing the maximum number of bindingCo_3_O_4_/PAn nanocomposites. Neither too large nor too small dilution ratio could be good for electric currency change, therefore, 1:400 was chose as the optimum dilution ratio.

The quantity of antigen immobilized on the surface of electrode could directly affect the sensitivity of the sensor, so the impact of artificial antigen concentration was investigated (Fig. 3C). With the increase of artificial antigen concentration, the change of electric current showed a decrease trend at the beginning, but reached a peak with the dilution ratio being 1:1600. When the dilution ratio was over 1:1600, the amount of immobilized antigen became too large and a larger steric hindrance was produced among antigen molecules, which reduced the ratio of antigen-antibody binding and further caused the reduction of the electronic signal change of the sensor. Hence, 1:1600 was chosen as the optimum dilution ratio.

Before indirect competitive immunoassay was performed, the response to azocylotin antibodies at different titers was investigated using the prepared probe in the samples which did not contain azocylotin (Fig. 3D). It showed that the electric current decreased with the increase of dilution ratio of azocylotin antibody, and the antibody was in super-saturation when the dilution ratio ranged from 1:100 to 1:200 However, if the solution was continuously diluted, the current response increased suddenly, and a peak value was achieved when the dilution ratio was 1:400, indicating that the increase of antibody concentration was proportional to binding rate if the dilution ratio was between 1:200 and 1:400. Moreover, it also revealed that the increase of antibody concentration had little effect on binding ratio if the dilution ratio was between 1:100 and 1:200, suggesting that the antigen might be in super-saturation. Therefore, 1:400 was considered as the optimum dilution ratio of azocyclotin antibody.

Antigens and antibodies are usually amphoteric substances with certain isoelectric points, and their spatial structures and the ionic station of groups are different under different pH values, which could influence the charging and affinity. The current responses of the immunosensor in was investigated at pH 4∼9 ((Fig. 3E). The current change of electrode of the sensor reached a peak value, and the enzyme activity was the best at pH=4.0. Therefore, PBS (1/15 M) at pH=4 was chose as the base solution.

Transforming immune signals into electrochemical signals through the enzyme catalysis of labeled HRP on the secondary antibody is a common method of electrochemical immunosensor. However, the signals generated by traditional single-labeled method are too weak to improve sensitivity of the sensor. The multi-labeled antibody was produced by using silver nanometric copper rod because gold nanorods have good biocompatibility and conductivity, and the surface effect of nanometric copper rod was good for absorbing more enzymes. A secondary antibody molecule could be linked to multiple enzyme molecules by using nano-materials. The system built by multi-labeled secondary antibody could obtain much larger signals than single-labeled system when the concentrations of samples are the same, thus, amplification of the effective signal is achieved to greatly improve the sensitivity of the sensor.

In order to explain the superiority of multi-labeled method, a set of control experiments were conducted using single-labeled and multi-labeled secondary antibodies, respectively, under the same conditions (Fig.4). It indicated that in the cyclic voltammogram, redox peak current change was 0.6 and 1.2 mA after adding 25 μl 0.48M H_2_O_2_ into the single-labeled and multi-labeled systems, respectively, suggesting that multi-labeled secondary antibodies truly have the capacity to amplify the signal of the enzyme and further improve the sensitivity of the sensors.

**Fig 4.**
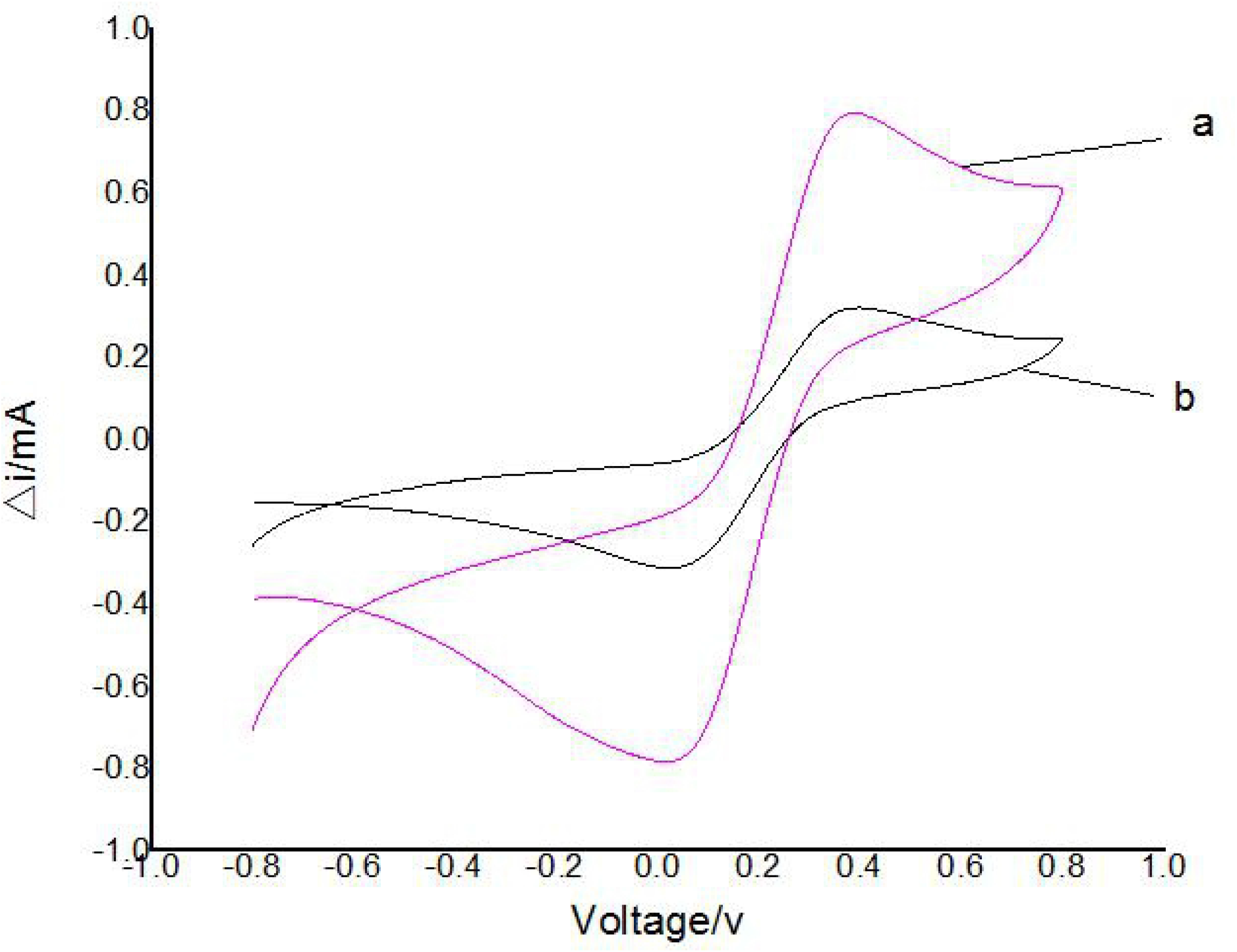
Contrastive analysis of multi-labeled secondary antibody and single-labeled secondary antibody. (a) Multi-labeled secondary antibody sensor, (b) single-labeled secondary antibody sensor.

The azocyclotin samples were diluted into a series of standard solutions with concentrations (0, 2, 4, 6, 8 and 10 μg/ml), and the standard curve was generated by indirect competitive assay by using PBS buffer as the control. As showed in the Fig. 3B, there was a significant liner relationship between frequency response and log of azocyclotin concentration ranging from 0 to 10μg/m1 [I(μA) =55.38X (μg/mL)+46.53]. The correlation coefficient was 0.9744, and the limit of detection (LLD) of the sensor was 0.01 μg/ml (Fig.5).

**Fig 5.**
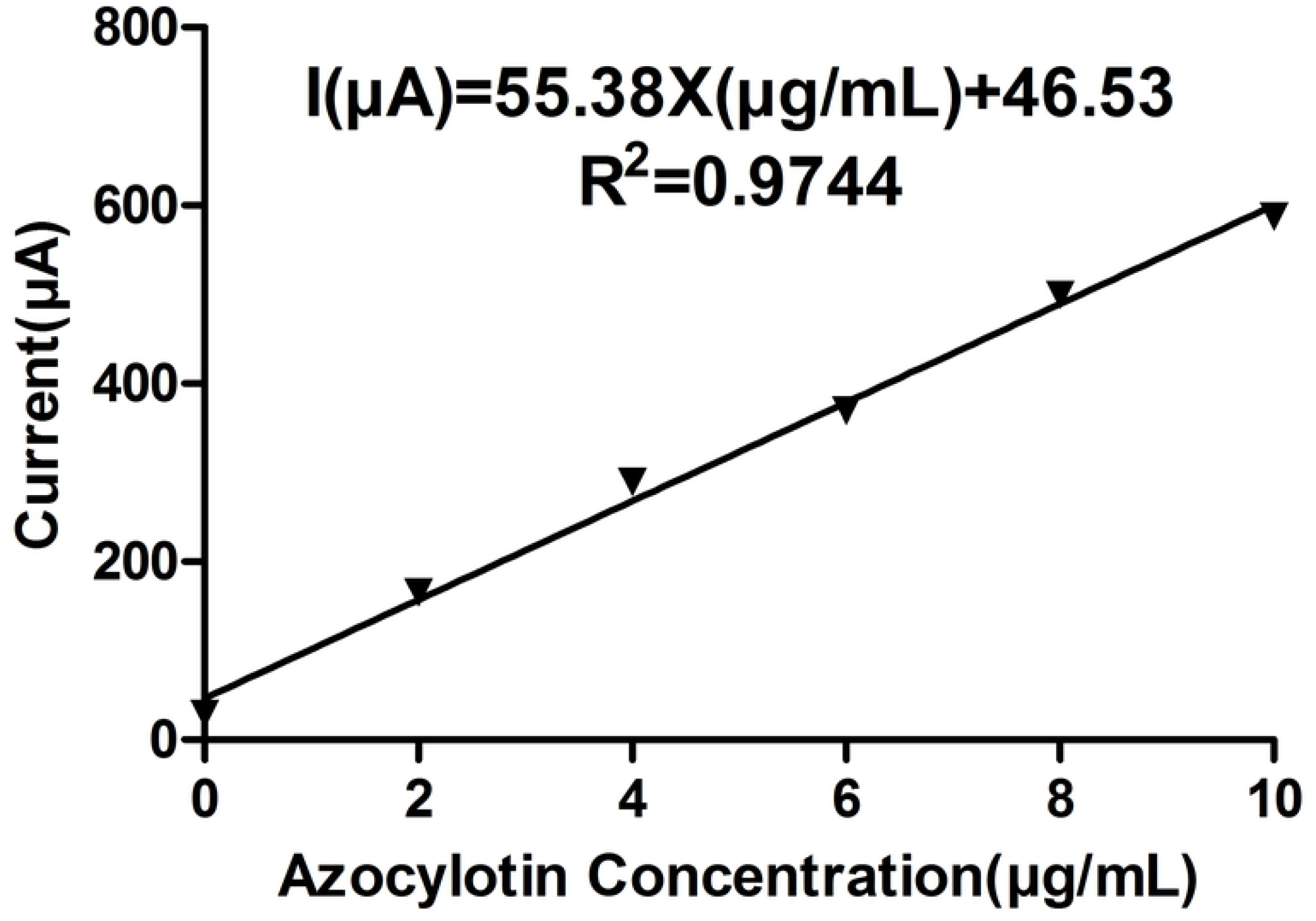
Standard curve of the determination results of azocylotin standard using Co_3_O_4_/PAn sensor.

### Accuracy and regeneration of electrochemical immunosensor

The formula: recovery rate= (measured value of spiked samples-measured value of the sample)/spiked sample amount × 100%, is used to evaluate the conformity between the measured value and real value. The adding recovery rates of apple and orange samples were shown in Table 1. The recovery rates were over 88% with variable coefficient less than 5% for the two agricultural products. The recovery rate over 100% might be caused by the experimental error or background. The background value of the samples had great impacts on the recovery rate of the reference standards with low concentration. Generally, if the sample had lower background value, there might be the higher the recovery error of the reference standards.

**Table 1.**
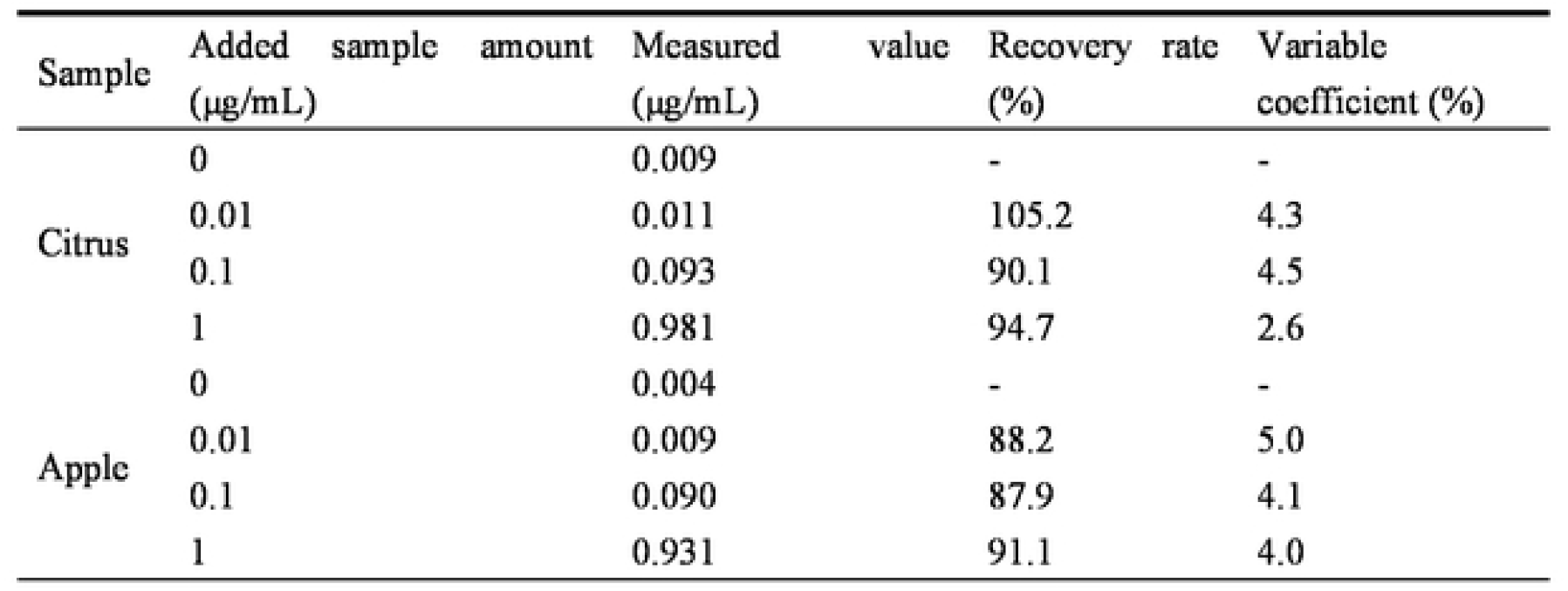
Added recovery rate of azocylotin in citrus and apple determined using Co_3_O_4_/PAn sensor

The recovery experiments were carried out for the produced immunoelectrodes. After detecting the samples of the standards, citrus and apple, the immunoelectrodes were washed 3 times, with acidic PBS buffer solution and then was used to test the standards again. The results were wonderful and had a good repeatability, indicating a better retrievability of the electrode.

## Discussion

In the current study, a kind of immune-electrochemistry sensor system based on the indirect competition method, which can be used to determine the residual azocylotin concentration in agricultural products, was constructed. The basic principle is that azocylotin artificial antigen coupled with BSA is coated on the surface of the working electrodes which were covered with a thin layer of Co_3_O_4_/PAn magnetic nanoparticles, and free azocylotin molecules in the solution are bound competitively to azocylotin monoclonal antibodies in the solution with azocylotin coupling antigens on the working electrodes. The immune reactions between antibodies and coupling antigens could be measured by voltammetry. The immunosensor was used to detect the pesticide residues of agricultural products like citrus and apple, and the sensitivity, accuracy and reproducibility were evaluated. Ranging from 0 to 10 μg/ml, the standard curve was as follows: I(μA) =55.38 x (μg/mL) + 46.53, with the correlation coefficient of 0.9744, with limit of detection of 0.01 μg/ml, the recovery rate of the standard in the relevant samples > 88% and variable coefficient < 5%. The generated electrodes still have reliable detection sensitivity after immersion cleaning with acidic PBS solution. In this experiment, the thin-layer electrodes, which were created using Co_3_O_4_/PAn magnetic nanoparticle has reliable performance, and can markedly increase the detection sensitivity to micromolecule pesticides such as azocylotin. Hence, it has a wonderful application value.

Electrochemical immunosensor, possessing the advantages of rapidness, high sensitivity and cheapness, can be widely applied in the medical, food, industrial and environmental area. In particular, by integrating with nanotechnology, the electrochemical immunosensor had become the top in the development and application of biosensor [15-17]. One objectives of making immunosensor based on thin-layer electrode is to improve the detection sensitivity and specificity to the analytes, and the other important objective is realizing the integration and miniaturization of electrochemistry detection system and sample separation system, thus achieving rapid, large-scale and on-site detection for pesticide residues in the samples [18,19]. For example, by combining micro-electrical mechanical system with micro-fluidic technique, the Co_3_O_4_/PAn thin-layer electrode and reference electrode were integrated into the pre-treatment system of the samples with the pesticide residues, so as to establish the laboratory-level rapid detection system of pesticide residues, which make the bulk of samples could be rapidly detected indoors [20]. Meanwhile, with the continuous integration of the micro electro mechanical system technology and nanotechnology into the sensor technology field, the electrochemical immunosensors tend to become microminiaturization and integration increasingly, and a variety of portable intelligentized electrochemical immunosensors will be gradually applied to disease diagnosis, food and environment monitoring [21,22]. With low cost, high sensitivity and good stability in electrochemical immunosensors, their technical progress will accelerate the commercial process in different fields.

## Acknowledgements

We would like to thank Dr Leilei Zhan for their helpful discussions and critical reading of the manuscript.

